# In the presence of population structure: From genomics to candidate genes underlying local adaptation

**DOI:** 10.1101/642306

**Authors:** Nicholas Price, Lua Lopez, Adrian E. Platts, Jesse R. Lasky, John K. McKay

## Abstract

Understanding the genomic signatures, genes, and traits underlying local adaptation of organisms to heterogeneous environments is of central importance to the field evolutionary biology. Mixed linear mrsodels that identify allele associations to environment, while controlling for genome-wide variation at other loci, have emerged as the method of choice when studying local adaptation. Despite their importance, it is unclear whether this approach performs better than identifying environmentally-associated SNPs without accounting for population structure. To examine this, we first use the mixed linear model GEMMA, and simple Spearman correlations, to identify SNPs showing significant associations to climate with and without accounting for population structure. Subsequently, using Italy and Sweden populations, we compare evidence of allele frequency differentiation (*F_ST_*), linkage disequilibrium (LD), fitness variation, and functional constraint, underlying these SNPs. Using a lenient cut-off for significance, we find that SNPs identified by both approaches, and SNPs uniquely identified by Spearman correlations, were enriched at sites showing genomic evidence of local adaptation and function but were limited across Quantitative Trait Loci (QTL) explaining fitness variation. SNPs uniquely identified by GEMMA, showed no direct or indirect evidence of local adaptation, and no enrichment along putative functional sites. Finally, SNPs that showed significantly high *F_ST_* and LD, were enriched along fitness QTL peaks and cis-regulatory/nonsynonymous sites showing significant functional constraint. Using these SNPs, we identify genes underlying fitness QTL, and genes linking flowering time to local adaptation. These include a regulator of abscisic-acid (*FLDH*) and flowering time genes *PIF3, FIO1*, and *COL5*.

## Introduction

Populations of a species may inhabit different environments where local selection pressures favor a combination of (multivariate) phenotypes (Leimu and Fischer 2008; Conover, et al. 2009; Hereford 2009; Savolainen, et al. 2013). Once locally adapted, the resident genotype is expected, on average, to have a higher relative fitness than a foreign genotype (Kawecki and Ebert 2004). Despite the widespread evidence of local adaptation in many taxa (Leimu and Fischer 2008; Jeong and Di Rienzo 2014; Arguello, et al. 2016), our understanding of the traits involved, its genetic basis, and its environmental underpinnings is still at an infant stage (Savolainen, et al. 2013; Tiffin and Ross-Ibarra 2014; Wadgymar, et al. 2017).

In a variety of species, reciprocal transplant and common garden/laboratory experiments have showed significant adaptive differentiation between natural populations inhabiting different environments (Via 1991; Hendry, et al. 2002; Savolainen, et al. 2007; Ågren and Schemske 2012; Kaufmann, et al. 2017; Phifer-Rixey, et al. 2018). Furthermore, in plants and animals, mapping experiments have uncovered Quantitative Trait Loci (QTL) for traits that are thought to underlie local adaptation (Colosimo, et al. 2004; Oakley, et al. 2014; Yang, et al. 2016; Ågren, et al. 2017), in addition to QTL explaining fitness differences across environments (Ågren, et al. 2013; Anderson, et al. 2013). Despite the importance of QTL studies in providing direct evidence for local adaptation (Ågren, et al. 2013), in many instances they provide a low resolution for its genetic basis, and in practical terms are time consuming, expensive, and labor intense (Joosen, et al. 2009).

With the advent of low-cost, and fast, next generation sequencing (Henson, et al. 2012), higher resolution population genomics approaches have emerged as the new means for examining the genetic basis of local adaptation. (Lachance and Tishkoff 2013; Savolainen, et al. 2013; Tiffin and Ross-Ibarra 2014; Sork 2017). In brief, these methods include: (a) identifying single nucleotide polymorphisms (SNPs) showing significant allele frequency differentiation between populations (*F_ST_*) (Beaumont and Balding 2004; Foll and Gaggiotti 2008; de Villemereuil and Gaggiotti 2015); (b) identifying genomic regions showing significant increases in linkage disequilibrium (Jacobs, et al. 2016) or composite likelihood ratios for recent sweeps (DeGiorgio, et al. 2016; Huber, et al. 2016); and (c) alleles showing significant correlations to environment/climate (Hancock, et al. 2011; Jones, et al. 2012; Lasky, et al. 2012; Lasky, et al. 2014; Pluess, et al. 2016; Yeaman, et al. 2016; Monroe, et al. 2018; Price, et al. 2018). The latter approach has gained particular attention because it can be implemented on the basis of individual (as opposed to population-based) sampling, and furthermore it provides a direct link to ecologically relevant factors (e.g., climate).

Despite the ability of population genomic methods to identify candidate genetic variation underlying local adaptation, it is hard to disentangle the effects of selection from demographic history (Lotterhos and Whitlock 2014; Hoban, et al. 2016). Geographically varying environments can generate population structure in the regions of genes involved in adaptation, even under conditions (e.g. high gene flow) that do not generate population structure at a genome-wide level (McKay and Latta 2002). The degree to which population structure influences the patterns of genome-wide LD and therefore the occurrence of false positives has emerged as a critical hurdle (Platt, et al. 2010). To limit the number of spurious associations, studies usually estimate population structure using different methods (Price, et al. 2010), incorporate population structure (Yu, et al. 2006; Kang, et al. 2010; Wang, et al. 2011; Zhou and Stephens 2012) and/or geographic structure (Lasky et al. 2012) into statistical models, and finally test whether certain loci explain significantly higher variation in environment/climate than population structure itself (Hancock, et al. 2011; Lasky, et al. 2012; Fischer, et al. 2013; Huber, et al. 2014; Lasky, et al. 2014; Monroe, et al. 2016; Rellstab, et al. 2017; Frachon, et al. 2018; Lasky, et al. 2018; Price, et al. 2018). While such approaches may limit the number of false positives, they can also lead to false negatives (Bergelson and Roux 2010; Anderson, et al. 2011). According to simulations (Forester, et al. 2018), when selection is spatially autocorrelated, accounting for population structure reduces the power to detect loci under selection (Forester, et al. 2018).

The negative effects of accounting for population structure, may explain the reduced signal of genetic convergence to climate among distantly related conifers (Yeaman, et al. 2016). Furthermore, in *Arabidopsis thaliana*, it may underlie the lack of significant associations between climate-correlated SNPs and fitness QTL exhibiting genetic tradeoffs (Price, et al. 2018); and the lack of SNPs showing significant associations to drought survival among Eurasian accessions (Exposito-Alonso, et al. 2018). Finally, in the plant *Capsella bursa-pastoris*, accounting for population structure explained all variation in gene expression among ecotypes that differed in important life-history traits such as flowering time and circadian rhythm (Kryvokhyzha, et al. 2016).

In conjunction to population genomic signatures of selection or significant associations to climate, genetic variation underlying local adaptation is expected to be enriched along sites that are functional and influence fitness. SNPs showing significant associations to climate were found to be significantly enriched among nonsynonymous, but also synonymous variation (Hancock, et al. 2011; Lasky, et al. 2012). The enrichment among synonymous variation (which largely evolve neutrally) maybe the result of linkage disequilibrium due to neutral processes but also background selection (Charlesworth, et al. 1993) and/or hitchhiking (Gillespie 2000). A stricter enrichment test will be one that controls for sequence conservation along coding and non-coding sites. Sites that are highly conserved among species (Miller, et al. 2007; Haudry, et al. 2013; Hupalo and Kern 2013), are assumed to be under functional/selective constraint and functionally important — that is, due to purifying selection the number of tolerated mutations is very limited (Graur 2016). Therefore, SNPs showing significant evidence of local adaptation across highly constraint sites are more likely to be true positives.

In the current study, we first examine how accounting for population structure in genome-wide associations to climate may affect our ability to provide a comprehensive picture on the genetic basis of local adaptation; and secondly, we use an approach that accounts for genetic signatures of selection and functional constraint to detect potential genes underlying local adaptation.

More specifically, during the first phase we use 875 *A. thaliana* Eurasian accessions and identify SNPs showing significant correlations to Minimum Temperature of Coldest Month (Min.Tmp.Cld.M) using a mixed linear model that accounts for population structure (Zhou and Stephens 2012) (GEMMA-Genome-wide Efficient Mixed Model Association) and simple Spearman correlations (Spearman 1987) that did not account for population structure (we mainly focused on Min.Tmp.Cld.M —a proxy to winter temperature—because of significant evidence linking cold acclimation to local adaptation in wild populations (Ågren and Schemske 2012; Oakley, et al. 2014; Gienapp, et al. 2017; Oakley, et al. 2018)). Using the two sets of SNPs, we first separated them into ones identified by both methods (referred to as “Common” hereafter), and ones uniquely identified by each approach (referred to as “GEMMA” and “Spearman” hereafter). Thereafter, we compared evidence of local adaptation and function underlying the three sets of SNPs. More specifically, using Italy and Sweden re-sequenced genomes, and QTL explaining fitness variation between these populations (Ågren, et al. 2013), we examined: (a) the level of allele frequency differentiation and linkage disequilibrium underlying SNPs showing significant associations to climate before and after accounting for population structure; (b) how these SNPs were distributed along LOD score peaks of 20 fitness QTLs (Ågren, et al. 2013); and (c) how they were distributed along nonsynonymous and cis-regulatory sites showing significant functional constraint among plants species of the Brassicaceae family (Haudry, et al. 2013).

During the second phase, we use SNPs showing significant evidence of local adaptation and function, to identify potential genes underlying fitness QTL and flowering time variation between Italy and Sweden populations. Flowering time is a life history trait that is thought to play a significant role in local adaptation to climate (Hall and Willis 2006; Verhoeven, et al. 2008; Sandring and Agren 2009; Dittmar, et al. 2014; Ågren, et al. 2017), and whose genetic basis has been thoroughly studied (Salomé, et al. 2011; Sasaki, et al. 2017). To re-examine evidence linking flowering time to climate adaptation we used the following data: (1) a list of genes that were experimentally shown to affect flowering time; (2) high confidence QTL explaining flowering time variation between Italy and Sweden populations (Ågren, et al. 2017), and (3) flowering time estimates for Arabidopsis Eurasian accessions (1001 Genomes Consortium 2016).

## Materials and Methods

### Detecting associations to climate with and without accounting for population structure

To compare allele associations to climate with and without accounting for population structure we focused on the climate variable Minimum Temperature of Coldest Month (Min.Tmp.Cld.M). Using a SNP genotype matrix for a panel of 1,135 globally distributed accessions downloaded from the 1001 Genomes database, we filtered out accessions from outside the native Eurasian and North African range of *A. thaliana*, as these accessions may have weaker patterns of local adaptation (Lasky, et al. 2012). We also filtered out accessions that were likely laboratory escapees or contaminants (Pisupati, et al. 2017), leaving 875 accessions. After we filtered for biallelic SNPs with minor allele frequency >0.05, we tested association with home climate of ecotype and tested for potential confounding effects of population structure using the software “gemma” (Zhou and Stephens 2012). The parameters used in gemma were a MAF of 0.05 (default 0.01) and a missingness threshold of 0.05. For the linear mixed model option, we used Wald test (default) to test for significant associations to climate. We tested models where home climate was a function of SNP allele, and the association p-values we report are for the null hypothesis that the mean climate occupied by the two alleles is equal (Lasky, et al. 2014). Using the same set of SNPs we estimated correlations to climate using simple “Spearman” correlations (Spearman 1987) and not accounting for population structure. To estimate p-values for the Spearman correlations we used the ‘cor.test’ function implemented in R (Team 2009).

### Estimates of selection and candidate functional variation underlying populations in North Sweden and South Italy

In addition to associations to climate, genomic signatures of selection were examined in North Sweden and South Italy populations that represent the most northern and southern tips of Eurasia. The accessions used, including latitude and longitude coordinates are found in the Supplementary data file. Evidence of local adaptation/selection across SNPs between Italy and Sweden populations was measured using absolute allele frequency differentiation (*F_ST≈_* |*f_N.Sweden_* – *f_S.Italy_*|) and linkage disequilibrium (LD) between a SNP and its neighboring SNPs with a 20kb window. LD was measured using the package ‘PLINK’ (Purcell, et al. 2007) and it was estimated as the mean square coefficient of correlation 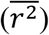. For evidence of recent sweeps we used previously calculated (Price, et al. 2018) Composite likelihood Ratios (CLR’s) that were computed using Sweepfinder2 (DeGiorgio, et al. 2016). We focused on CLR’s in North Sweden because in other populations signals identified were very weak (Long, et al. 2013; Huber, et al. 2014; Price, et al. 2018).

To narrow down SNPs to ones that are more likely to underlie differences in function/expression of protein-coding genes we focused on cis-regulatory and nonsynonymous variation that was found along sites showing significant functional constraint. We regarded cis-regulatory SNPs as those found within 1 kb upstream from the transcriptional start site of a gene (Zou, et al. 2011; Pass, et al. 2017), unless these sites were found in transcribed regions of other genes (in which case they were excluded). To call nonsynonymous variation we used bi-allelic sites, we used a publicly available python script (callSynNonSyn.py; archived at https://github.com/kern-lab/), and gene models downloaded from the TAIR database (TAR10 genome release) (Berardini, et al. 2015). To annotate regions showing significant functional constraint across the *A. thaliana* genome we used phastCons scores (Siepel, et al. 2005) derived using a nine-way alignment of Brassicaceae species from the study by Haudry et al. (2013). We defined conserved regions as those with a score >=0.8 over blocks of >=10 nucleotides.

### Fitness and flowering time QTL underlying Italy and Sweden populations

Quantitative trait loci explaining fitness variation between natural *A. thaliana* Italy and Sweden populations were retrieved by the study of Ågren et al. (2013). These 20 fitness QTL were assembled into 6 genetic tradeoff QTL (Ågren, et al. 2013), however we treated them as independent given the very long genetic distances between fitness QTL peaks (Supplementary data). Furthermore, we retrieved high confidence QTL explaining flowering time variation between these populations (Ågren, et al. 2017) (Supplementary data).

### Circular permutation tests

To test whether SNPs showing significant correlations to Min.Tmp.Cld.M and/or SNPs showing high *F_ST_* and LD between Italy and Sweden populations are enriched along QTL peaks and among cis-regulatory/nonsynonymous variation at sites showing significant functional constraint we used a 1,000 circular permutations (Fig. 1).

**Fig. 1.**
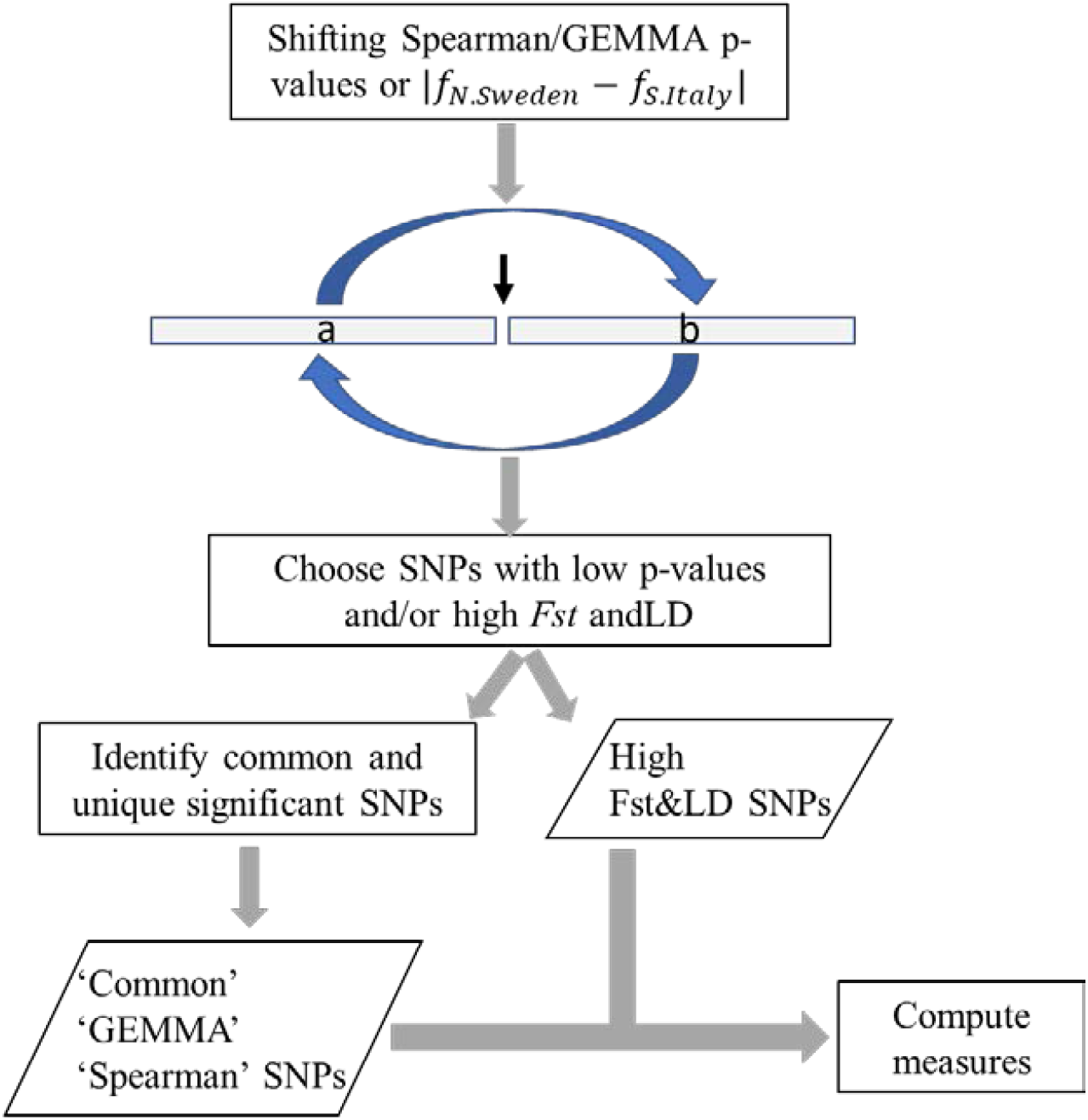
Diagram of circular permutations used to build null distributions for various measures. The first step involves choosing a random location along the genome and shift p-values or allele frequency differentiations. In the next step, when examining climate associations, we chose SNPs with a p-value less than a specified threshold. We also samples of SNPs showing a high *F_ST_* and LD. SNPs showing “significant” correlations to climate were further partitioned into those showing “significance” using both Spearman and GEMMA correlations (“Common”) or were unique to each approach (“GEMMA” or “Spearman”). Using the final set of SNPs, we computed various measures of interest.

Fig. 1 depicts the steps involved in the circular permutations. In brief, during the first step p-values associated with climate correlations or allele frequency differentiations were shifted based on a random SNP along the genome. The second step involved choosing SNPs showing “significant” associations to climate (p-value < 1^st^ percentile of Spearman/GEMMA p-value distributions), or high allele frequency differentiations and LD (|*f_N.Sweden_* – *f_S.Italy_*|>0.70 and LD>0.19: 0.70 and 0.19 represent the 95^th^ percentiles of the distributions). For climate associations, SNPs were partitioned into those showing “significance” using both Spearman and GEMMA associations (“Common”) or those showing “significance” using only one approach (“GEMMA” or “Spearman”).

The final sets of SNPs were used to estimate the following measures: (a): % of SNPs found within a certain distance (100, 200, ….600 kb) upstream and downstream of the 20 fitness QTL peaks; and (b) the proportion of cis-regulatory and nonsynonymous SNPs that were within regions showing significant functional constraint.

### Sliding window analysis of chromosomal variation SNPs showing evidence of local adaptation

To detect chromosomal regions with a high proportion of SNPs showing significant evidence of local adaptation we used a sliding window approach. Specifically, for a window size of 20kb and a step size of 1kb we estimated the ratio of SNPs showing a specific requirement (e.g., *f_N.Sweden_* – *f_S.Italy_*|>0.70 and LD>0.19) over the total number of SNPs within a 20kb window.

### Flowering time estimates for A. thaliana Eurasian accessions and candidate flowering time genes

Estimates of flowering time for the 835 Eurasian *A. thaliana* accessions were downloaded from the study by Alonso-Blanco et. al (2016) (1001 Genomes Consortium 2016). In brief, plants were grown in growth chambers with the following settings: after 6 days of stratification in the dark at 4°C, constant temperature of 16°C with 16 hours light / 8 hours darkness, 65% humidity. Flowering time was scored as days until first open flower. See Alonso-Blanco et al. (2016) for further details. A set of genes that were experimentally verified to affect flowering time was downloaded from Prof. Dr. George Coupland website (https://www.mpipz.mpg.de/14637/Arabidopsis_flowering_genes)

### Constructing rooted gene trees

To build neighbor joining trees of genes showing significant local adaptation we downloaded 1:1 orthologs between *Arabidopsis thaliana* and outgroups *Arabidopsis lyrata* and *Capsella rubella* from the Phytozome database (Goodstein, et al. 2012) and after aligning the coding sequences with MAFFT (Katoh and Toh 2008) we used MEGA (Tamura, et al. 2013) to build a rooted gene trees.

## Results

### Genomic signatures of local adaptation and selection captured by climate associations in Italy and Sweden populations

Genome wide correlations to Minimum Temperature of Coldest Month (Min.Tmp.Cld.M) were examined using simple Spearman correlations (Spearman 1987) and a mixed model called GEMMA (Zhou and Stephens 2012) that accounted for putative population structure. Fig. S1 depicts the p-value distributions that were obtained when testing for significance using GEMMA and Spearman correlations. P-values underlying Spearman correlation were skewed towards very low p-values; many of which are likely false positives. Nonetheless, we compared the results obtained by the two methods using different percentiles of the p-value distributions as cutoffs for significance (10^th^, 5^th^, 1^st^ percentiles). For significant SNPs that were segregating between North Sweden and South Italy populations the percent of SNPs that were identified by both approaches (“Common”) at the three significance levels was ~20% (Fig. S2).

When comparing the average absolute allele frequency differentiation |*f_N.Sweden_* – *f_S.Italy_*| and LD 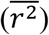 across SNPs that were unique to each approach (“Spearman” or “GEMMA”) and SNPs identified by both methods (“Common”) the set of “Common” and “Spearman” SNPs showed significantly higher allele frequency differentiation and LD than the genome average (Fig 2A-2B). On the other hand, SNPs that were unique to “GEMMA” did not show any large differences in *F_ST_* and LD from the genome average (Figs 2A-2B). Furthermore, across “Common” and “Spearman” SNPs we see a decrease in (|*f_N.Sweden_* – *f_S.Italy_*|) and LD as the threshold of significance becomes less stringent (Figs 2A-2B). This is indicative that stronger associations capture SNPs showing stronger evidence of local adaptation and recent selection. Therefore, for any further tests we used the set of SNPs that were significant using the 1^st^ percentile of the p-value distributions as the cutoff.

**Fig. 2.**
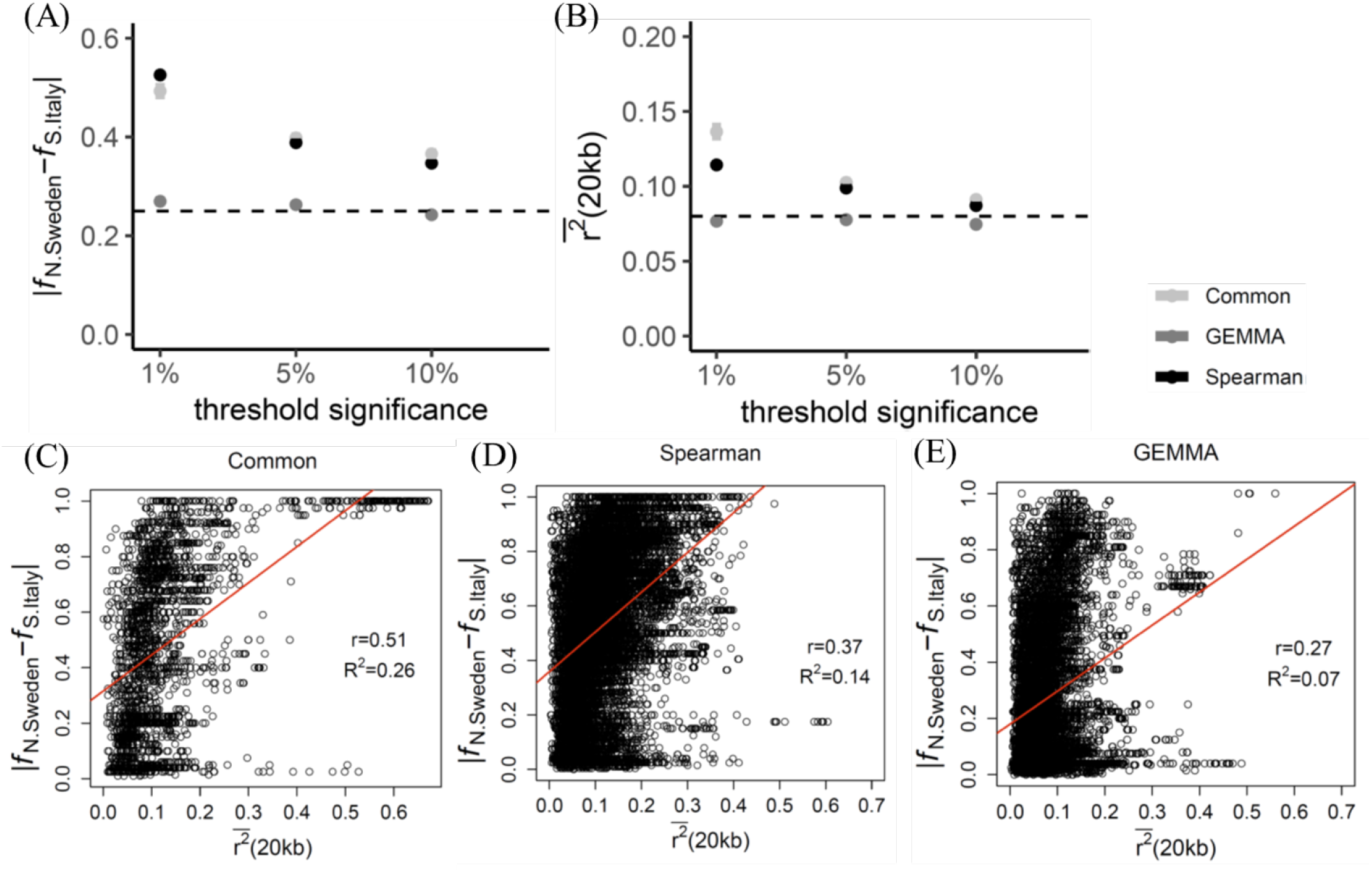
Comparing allele frequency differentiation and linkage disequilibrium across SNPs showing significant associations to Min.Tmp.Cld.M. (A-B) Average absolute allele frequency differentiation (|*f_N.Sweden_* – *f_S.Italy_*|) and linkage disequilibrium (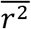) was significantly higher the genome average (dotted lines) across SNPs identified by both Spearman and GEMMA correlations (“Common”) and SNPs uniquely identified by Spearman correlations (“Spearman”). These sets of SNPs also showed a decrease in |*f_N.Sweden_* – *f_S.Italy_*| and 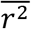 decreased as the the threshold for significance became more lenient (1→10 %) (95% CI’s which are not visible were estimated using a bootstrap approach). (C-E) A positive association between |*f_N.Sweden_* – *f_S.Italy_*| and 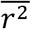 across the three sets of SNPs showing significant correlations to Min.Tmp.Cld.M when using the 1^st^ percentile of the p-value distributions (Fig. S1) as the significance level.

Local adaptation is expected to lead to an increase in allele frequency differentiation and LD at and near the site under selection; in other words, we would expect a positive correlation between |*f_N.Sweden_* – *f_S.Italy_*| and LD. As shown in Figs 2C-2E all three sets of SNPs showed significant associations between |*f_N.Sweden_* – *f_S.Italy_*| and LD. On the other hand, while “Spearman” SNPs were enriched in SNPs showing a |*f_N.Sweden_* – *f_S.Italy_*| near fixation (>0.90) the level of LD was lower (Fig. 2D) (also reflected in Fig. 2B). Finally, as expected given the small deviations from the genome average (Figs 2A-2B), “GEMMA” SNPs showed the poorest associations with a small sample of SNPs showing a |*f_N.Sweden_* – *f_S.Italy_*| between 0.60-0.80 and LD between 0.30-0.40 (Fig. 2E). Taken together, indicate indicate that “Common” and “Spearman” SNPs capture higher population genomic evidence of local adaptation, than SNPs uniquely identified by GEMMA.

### Detecting recent selection underlying fitness QTL is significantly reduced when accounting for population structure in GWA to climate

Direct evidence of local adaptation was shown in a study by Ågren et al. (2013) in which they identified 20 QTL explaining fitness variation between Italy and Sweden populations (Supplementary data). To examine population genomic evidence of local adaptation underlying these QTL we first examined |*f_N.Sweden_* – *f_S.Italy_*| and LD which are measures specific to Italy and Sweden populations. If the peaks of the QTL identified, are on average, near the site under selection we would expect a significant enrichment of high *F_ST_* and LD SNPs. Using different window sizes (100, 200, ….600 kb) from the QTL peaks we compared the observed proportion of high *F_ST_* SNPs (|*f_N.Sweden_* – *f_S.Italy_*| >0.70 —0.70 represents the 95^th^ percentile of the distribution), and the proportion of high *F_ST_* and LD SNPs (LD>0.19 — 0.19 represents the 95^th^ percentile of the distribution) to the expected proportion derived using a circular permutation test. As expected, we observed a significantly high proportion of high *F_ST_* (>*0.70*) SNPs and high *F_ST_* and high LD SNPs (Figs 3A-3B). A significant proportion was also observed at 200 kb, but as window sizes became larger the proportions became insignificant (<95^th^ percentile) and normalized to the genome average.

**Fig. 3.**
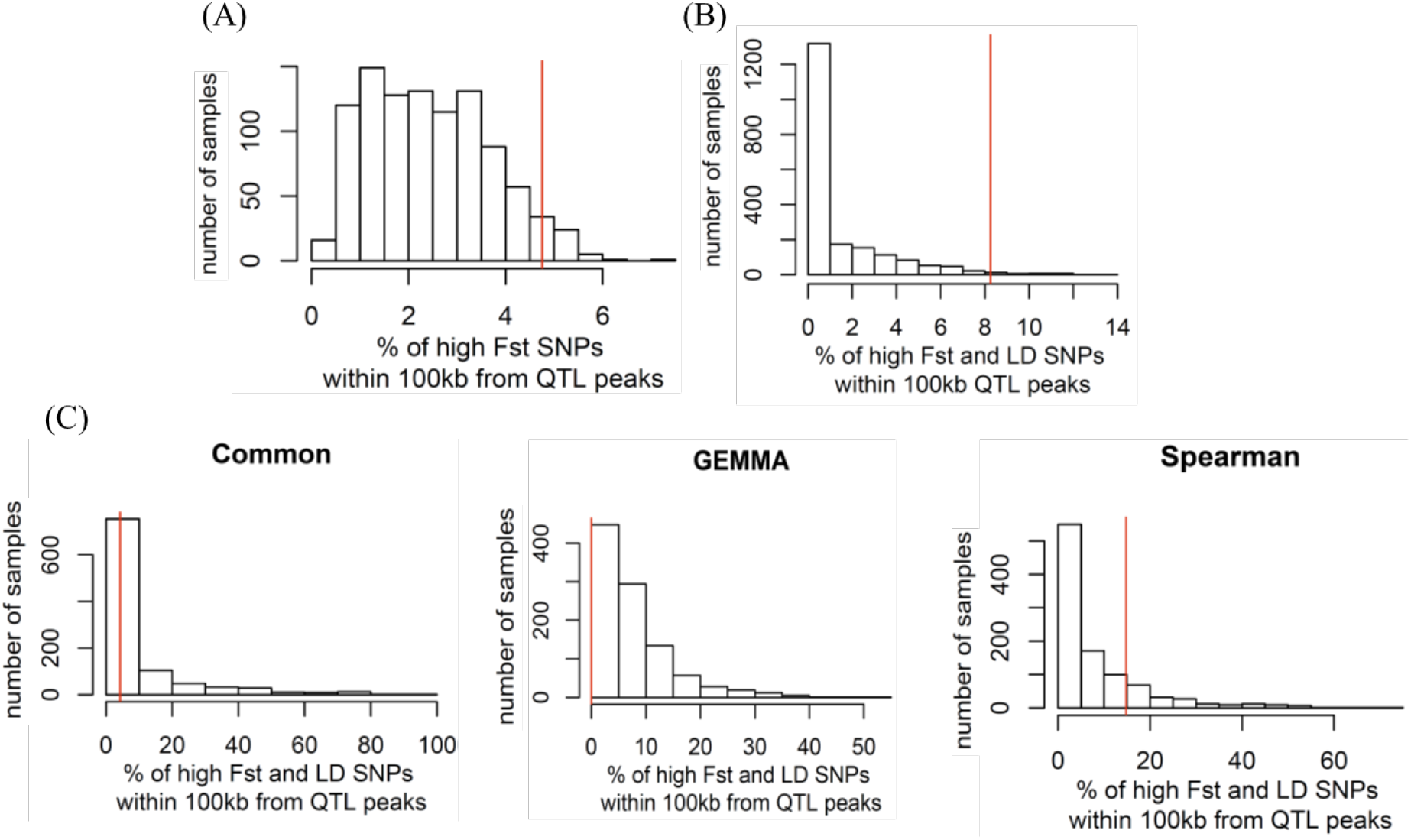
Distribution of SNPs showing population genomic evidence of local adaptation along fitness QTL peaks. (A) The observed proportion of SNPs showing significantly high allele frequency differentiation (|*f_N.Sweden_* – *f_S.Italy_*| > 0.70) (red line) in comparison to a permutation distribution of random proportions. The observed proportion was greater than the 95^th^ percentile. (B) The observed proportion of SNPs showing a |*f_N.Sweden_* – *f_S.Italy_*| > 0.70 and significantly high linkage disequilibrium 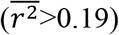, was also greater than the 95^th^ percentile of random proportions. (C) The observed proportions of SNPs showing significant associations to climate, in addition to a |*f_N.Sweden_* – *f_S.Italy_*| >0.70 and a 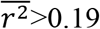. The proportion of “Common” and “GEMMA” SNPs near QTL peaks was very low, while the proportion “Spearman” SNPs was greater than the 80^th^ percentile of the distribution but less than the 95^th^.

Next, we performed the same test using SNPs showing significant associations to climate, and SNPs that showing significant associations to climate and a high *F_ST_* and high LD. When considering only associations to climate (Fig. S3) we find an enrichment of “Common” and “Spearman” SNPs (Fig. S3), but not “GEMMA” SNPs. On the other hand, when we also considered high *F_ST_* and high LD, none of the SNPs showed an enrichment (Fig. 3C). Among the three sets, Spearman SNPs showed the highest observed proportion (>80^th^ percentile). To determine whether the significantly lower proportion of “Common” SNPs showing a high *F_ST_* and high LD was due to the stringency of our cutoff, we calculated the average *F_ST_* and LD for SNPs with an |*f_N.Sweden_* – *f_S.Italy_*| and <0.70 and 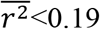. The average |*f_N.Sweden_* – *f_S.Italy_*| (≈0.26) and LD (≈0.06) were near or lower than the genome average (Fig. 2A-2B) to be considered as strong candidates underlying local adaptation.

Given that “Common” SNPs did not show an enrichment within 100kb of the 20 fitness QTL peaks examined (Supplementary data), we searched whether any of these were within the six tradeoff QTL that each span a large genomic region (Supplementary data). As shown in Fig. S4, most of the high *F_ST_* and high LD “Common” SNPs (181/209) were found within a 60kb region (14,719-14,781Mb) of a single genetic tradeoff (GT) QTL (GT QTL 2:2) (Fig. S4). Apart from being limited to a single GT QTL, given the multiple fitness QTL within GT QTL 2:2 (which spanned a length of ~4Mb); the multiple regions showing significant recent sweeps in Sweden; and the large number of windows showing a high proportion of high *F_ST_* and LD SNPs; it is highly unlikely that the “Common” set of SNPs cover all the causative genetic variation within GT QTL 2:2.

To further test for enrichment of “GEMMA” SNPs showing significant associations to climate and within 100 kb of fitness QTL peaks, we used an additional three climate variables (Table S1). Using a lenient cutoff for significance (FDR<0.1), we only identified a few additional SNPs within 100 kb of fitness QTL.

All in all, our results indicate that accounting for population structure when performing GWA to climate significantly reduces our ability to capture recent selection underlying fitness QTL.

### All but “GEMMA” SNPs show enrichment at cis-regulatory and nonsynonymous sites showing significant functional constraint

SNPs underlying local adaptation are expected to be significantly enriched at sites that affect function and/or expression of protein coding genes. Using the three sets of SNPs showing significant associations to climate (“Common”, “GEMMA”, and “Spearman”) and SNPs showing high *F_ST_* and LD (abbreviated as “*F_ST_*&LD”) we examined their distribution among cis-regulatory and nonsynonymous SNPs at sites showing significant functional constraint among Brassicaceae plants (Haudry, et al. 2013) (phastCons>0.8).

As shown in Figs 4A & 4B, the proportions of nonsynonymous/cis-regulatory “Common”, “Spearman”, and high “*F_ST_*&LD” SNPs, was significantly higher than expected by chance. On the other hand, “GEMMA” SNPs did not show a significant enrichment in any of the categories. When considering all the results obtained so far (Figs 2-4), unique “Spearman” SNPs seem to capture additional genetic variation underlying local adaptation (i.e., in addition to “Common” SNPs), while unique “GEMMA” SNPs do not.

**Fig. 4.**
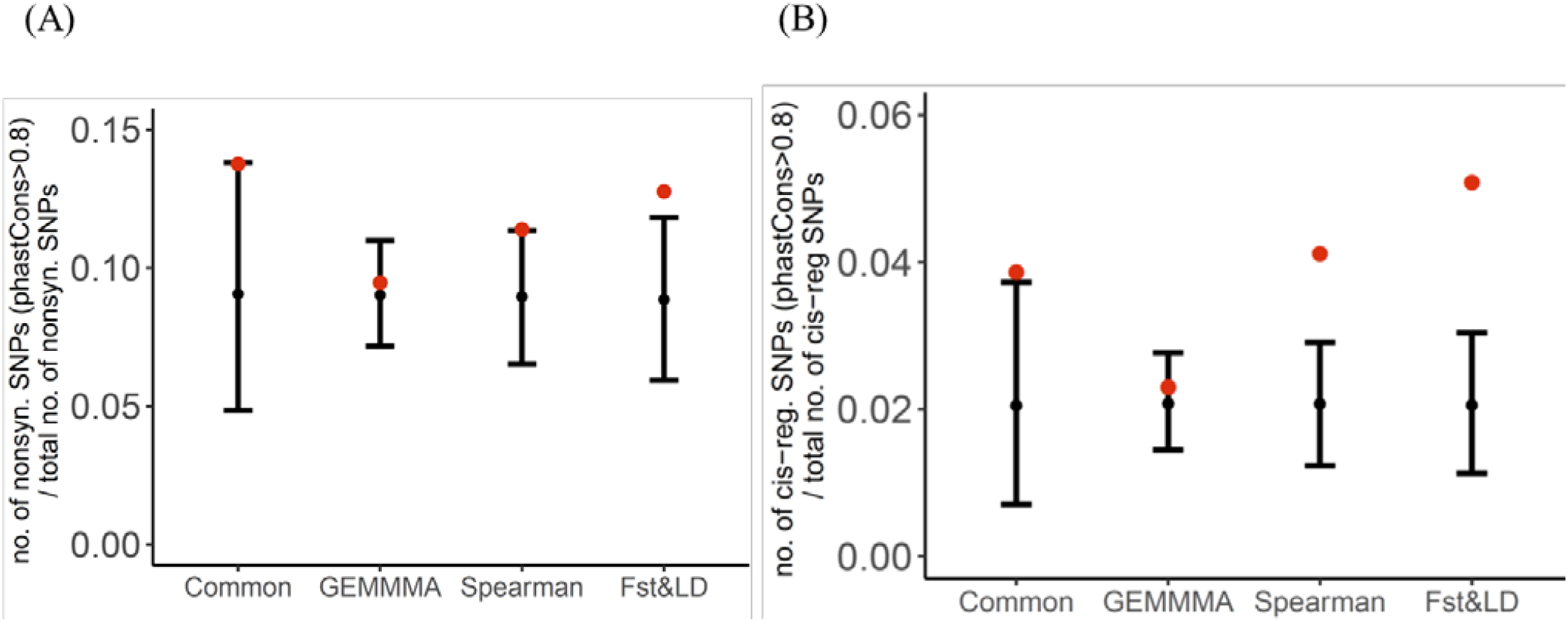
Proportion of cis-regulatory and nonsynonymous SNPs showing population genomic evidence of local adaptation and function (A) Red dots indicate the observed proportion of nonsynonymous SNPs showing associations to climate (“Common”, “GEMMA”, “Spearman”) or high *F_ST_* and LD (“*F_ST_*&LD”) and within coding regions showing significant functional constraint (phastCons>0.8). Expectations and 95% CI’s were derived using circular permutations (B) The observed proportions of cis-regulatory SNPs showing population genomic evidence of local adaptation and found within conserved regions.

### Flowering time genes showing significant evidence of local adaptation and underlying flowering time QTL

Given the significant enrichment of high *F_ST_* and LD SNPs, along QTL peaks (Fig. 3B) and conserved cis-regulatory and nonsynonymous sites (Figs 4A & 4B), we used them to detect potential genes that may underlie fitness QTL (note: we only focused on variants within 100kb of their peaks), in addition to examining evidence of local adaptation underlying a list of ~170 genes affecting flowering time (Supplementary data). To further narrow down on SNPs that are more likely to underlie the fitness QTL examined, we only considered variation that segregated between the parents used to derive the RIL’s (Ågren, et al. 2013). SNPs between the parental genomes were called in a previous study (Price, et al. 2018).

Our analysis resulted in 25 genes within 100 kb of fitness QTL peaks and spanning three genetic tradeoff QTL (2:2, 4:2, 5:5) (Table S2). Many of these were involved in interesting biological processes such as: response to different abiotic stress factors and the abscisic-acid signaling pathway which is important in abiotic stress response (Tuteja 2007) (Table S2). Among these genes, two of them (AT4G33360 (*FLDH*), AT4G33470 (*HDA14*)) showed strong expression GxE interactions (GxE interactions were identified in a previous study (Price, et al. 2018)) when Italy and Sweden plants were grown under cold acclimation conditions (4 °C) for two weeks (Gehan, et al. 2015). Interestingly, *FLDH* is a negative regulator of the abscisic acid signaling pathway (Bhandari, et al. 2010). As shown in Fig. 5A this gene was within a region of a genetic tradeoff QTL that showed a significantly high proportion of high *F_ST_* and high LD SNPs. Expression of *FLDH* under control and cold acclimation conditions was significantly lower in Sweden than Italy plants (Fig. 5A).

**Fig. 5.**
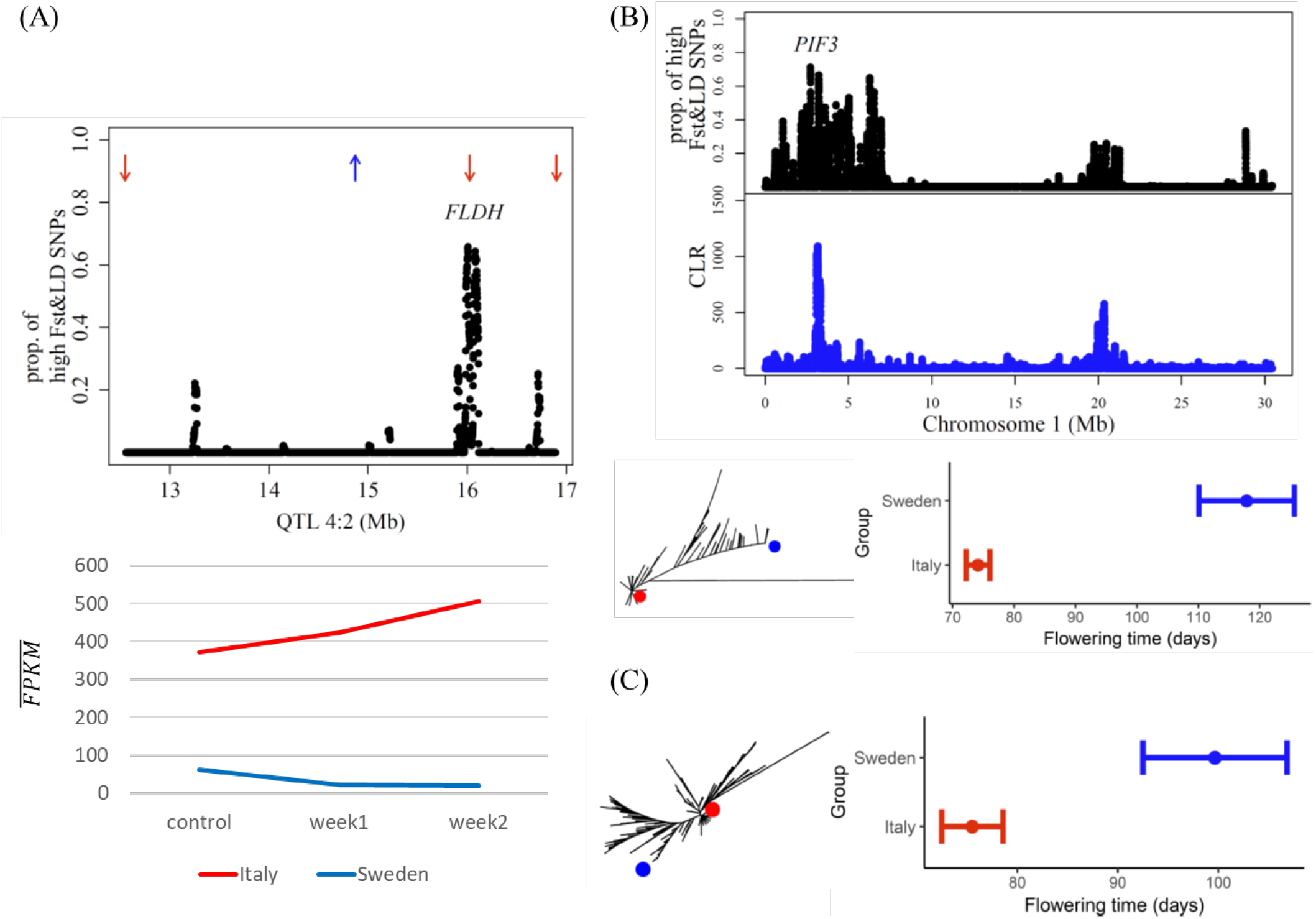
Fitness QTL and flowering time genes showing significant evidence of local adaptation along conserved cis-regulatory and nonsynonymous sites. (A) *FLDH* a negative regulator of ABA (Bhandari, et al. 2010) was found within a genetic tradeoff QTL 4:2 and 100 kb from a fitness QTL peak (red and blue arrows represent QTL where the Sweden genotype had lower fitness in Italy and higher fitness in Sweden, respectively). *FLDH* was found within a region showing a high proportion of high *F_ST_* and high LD SNPs. In Italy and Sweden plants it showed strong expression GxE interactions under control and cold acclimation conditions for two weeks (FPKM: Fragments Per Kilobase Million). (B) *PIF3* is a phytochrome interacting factor that has been found to affect flowering time (Oda, et al. 2004) and was found underlying a region along chromosome 1 that showed the largest Composite Likelihood Ratio (CLR) of a recent sweep in Sweden and windows with a high proportion of high *F_ST_* and high LD SNPs. A rooted phylogeny of the *PIF3* coding region indicated that Eurasian accessions sharing the same allele as the Sweden parent (blue dot) show significantly higher flowering time than accessions sharing the same allele as the Italy parent (red dot). (C) *COL5* is another gene that has been found to affect flowering time (Hassidim, et al. 2009) and in which Eurasian accessions show significant genetic differentiation and segregation in flowering time. This gene is also found within previously identified flowering time QTL (FlrT-5:4) (Ågren, et al. 2017) in which the Sweden genotype was associated with longer flowering time in both Italy and Sweden

Among the set of flowering time genes, we identified three (AT1G09530 (*PIF3*), AT2G21070 (*FIO1*), and AT5G57660 (*COL5*)) that contained high *F_ST_* and LD nonsynonymous SNPs within conserved coding regions. Among the three genes, *PIF3* was found along a chromosomal region that showed the highest CLR for a recent sweep in Sweden and a high density of SNPs showing high *F_ST_* and high LD (Fig. 5B).

Eurasian accessions sharing a similar allele as the Sweden parent showed longer flowering time than accessions sharing the same allele as the Italy parent (Fig. 5B). The same pattern was observed when examining *COL5* (Fig. 5C), a flowering time gene which was also found within a flowering time QTL (FlrT-5:4, Supplementary data). According to FlrT-5:4, the Sweden genotype was associated with longer flowering time in both Italy and Sweden (Ågren, et al. 2017). In conjunction, with its overlap to a genetic tradeoff QTL (Ågren, et al. 2017), indicates a possible role in fitness tradeoffs. Studies have attributed flowering time variation within FlrT-5:4 to *VIN3* (1001 Genomes Consortium 2016; Ågren, et al. 2017). Although it may be an additional candidate we did not find any significant genetic differentiation and selection along coding and cis-regulatory sites of *VIN3*.

When examining flowering time genes with high *F_ST_* and LD along cis-regulatory/nonsynonymous sites that did not show significant functional constraint we identified an additional nine genes; four of which were found within flowering time QTL (**FlrT**): AT1G14920 (GAI); AT1G53090 (SPA4); AT2G22540 (SVP); AT2G47700 (RFI2); AT4G32980 (ATH1-**FlrT4:1**); AT5G24470 (PRR5-**FlrT5:2**); AT5G62640 (ELF5); AT5G65050 (MAF2-**FlrT5:5**); and AT5G65060 (MAF3-**FlrT5:5**). ATH1 was found in genetic tradeoff QTL 4:2, while genes MAF2, and MAF3 were found within genetic tradeoff QTL 5:5 and within 100 kb of fitness QTL peaks.

## Discussion

In the quest to study the genetic basis of local adaptation using genome wide associations to environment, linear mixed models have emerged as a powerful tool given their ability to account for population structure while testing for significant associations (Yu, et al. 2006; Kang, et al. 2008; Kang, et al. 2010; Zhou and Stephens 2012). Although they provide a robust statistical framework for removing many false positives, the current study shows that such an approach may significantly limit our ability to understand the polygenic basis of local adaptation.

SNPs showing significant associations to climate after accounting for population structure, were not in line with high *F_ST_* and LD SNPs that showed a significant enrichment along fitness QTL peaks and coding/noncoding sites under functional constraint. Since these QTL are of large effect (Ågren, et al. 2013), the lack of significant associations is less likely to be a result of many alleles having a small effect on fitness. A more likely explanation is that accounting for population structure using genetic relatedness estimated from genome-wide SNPs, can lead to a significant number of false negatives. *Arabidopsis thaliana* however, is a simple, highly inbred species —to correctly assess the impact of accounting for population structure when examining the genetic basis of local adaptation there needs to be examination of other species with more complex life-history traits and evolutionary dynamics.

A large portion of SNPs that showed significant associations to climate, and SNPs that showed high *F_ST_* and LD between Italy and Sweden populations were enrichment among nonsynonymous/cis-regulatory variation at sites showing significant functional constraint. These results, further support an important role of cis-regulatory (Lasky, et al. 2014; Siepel and Arbiza 2014; Li and Fay 2017; Price, et al. 2018; Sackton, et al. 2019) and nonsynonymous variation (Nachman, et al. 2003; Coop, et al. 2009; Lasky, et al. 2012; Huber, et al. 2014; Svetec, et al. 2016; Price, et al. 2018) in adaptation. Among the list of candidate genes underlying fitness QTL and showing significant evidence of local adaptation at functionally constraint sites, we identified *FLDH. FLDH* is a negative regulator of abscisic-acid signaling (Bhandari, et al. 2010), that showed strong GxE interactions between Italy and Sweden plants under cold acclimation conditions. Abscisic-acid signaling is known to play an important role in abiotic stress response (Tuteja 2007), with many studies supporting its role in local adaptation to climate (Keller, et al. 2012; Lasky, et al. 2014; Kalladan, et al. 2017; Ristova, et al. 2017).

In addition to abscisic-acid signaling, our results provide further support for the important role of flowering time in local adaptation to climate. Among a list of genes that were experimentally shown to affect flowering time, we identified three genes (*PIF3*, *FIO1*, and *COL5*) that showed significant evidence of local adaptation and functional constraint along nonsynonymous sites. *FIO1* was previously shown to contain SNPs that showed significant associations to flowering time among natural Swedish lines (Sasaki, et al. 2015) and *COL5* was located within a QTL that explains flowering time variation among Sweden and Italy recombinant inbred lines (Ågren, et al. 2017). Finally, *PIF3*, a transcription factor that interacts with phytochromes (Soy, et al. 2012), has been implicated in multiple biological processes including early hypocotyl growth (Monte, et al. 2004), photomorphogenesis (Dong, et al. 2017), flowering time (Oda, et al. 2004), and regulation of physiological responses to temperature (Jiang, et al. 2017). This highly conserved transcription factor was found within a large region that showed significant evidence of local adaptation. This chromosomal region may involve a single causative variant, or a group of linked genes that interact with *PIF3 and* were under selection because they contributed to building an advantageous phenotype (Barton and Bengtsson 1986; Yeaman and Whitlock 2011).

When ignoring functional constraint, we identify a list of addition flowering time genes showing significant evidence of local adaptation along nonsynonymous/cis-regulatory sites. Genes such as SVP, MAF2, and MAF3 were previously associated with flowering time variation among natural *Arabidopsis* accessions (Caicedo, et al. 2009; Sasaki, et al. 2015). Although adaptation may involve sites that are not deeply rooted and/or under strong functional constraint, including additional plant genomes when estimating sequence conservation across species may increase our power to detect functionally important regions. As shown by studies examining adaptation in species ranging from bacteria (Maddamsetti, et al. 2017) to birds (Sackton, et al. 2019), addressing functional constraint can improve our understanding of its genetic basis.

Finally, the current study shows that we need a new statistical framework to examine genome wide associations to environment, and furthermore, it increases our understanding on the genes and traits that may underlie local adaptation in *A. thaliana*.

## Notes

#### Summary of Updates

modified Figure

