## Supplementary information for "In the presence of population structure: From genomics to candidate genes underlying local adaptation"

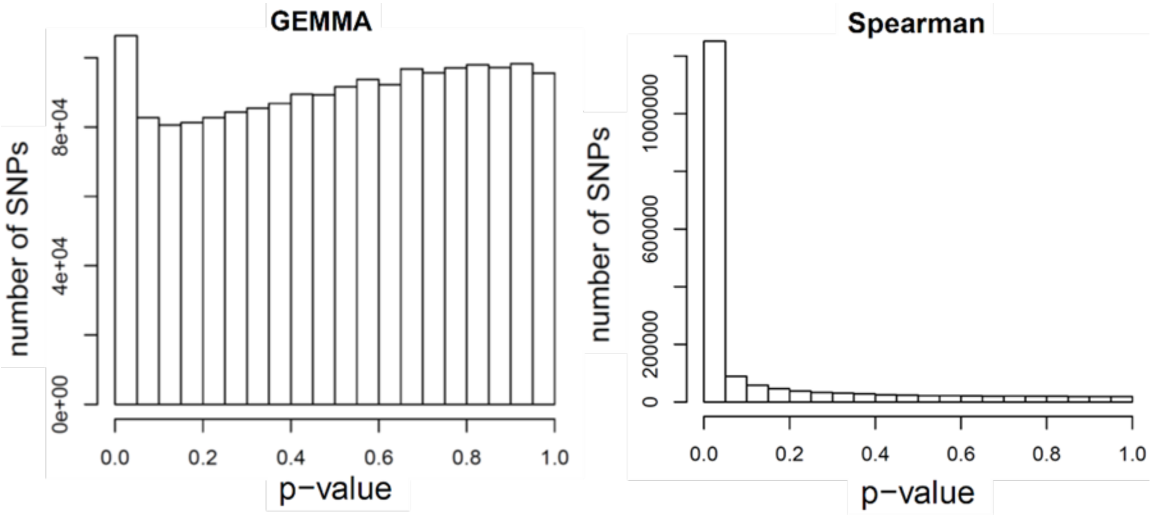


**Fig. S1**. The p-value distributions underlying associations to Minimum Temperature of Coldest Month (Min.Tmp.Cld.M) with and without accounting for population structure. The left figure depicts p-values that were obtained when testing for associations  to Min.Tmp.Cld.M using a mixed linear model that accounts for population structure and is implemented in GEMMA (Zhou and Stephens 2012). The right figure depicts p-values obtained when testing for associations to Min.Tmp.Cld.M using simple Spearman correlations that did not account for population strcture.

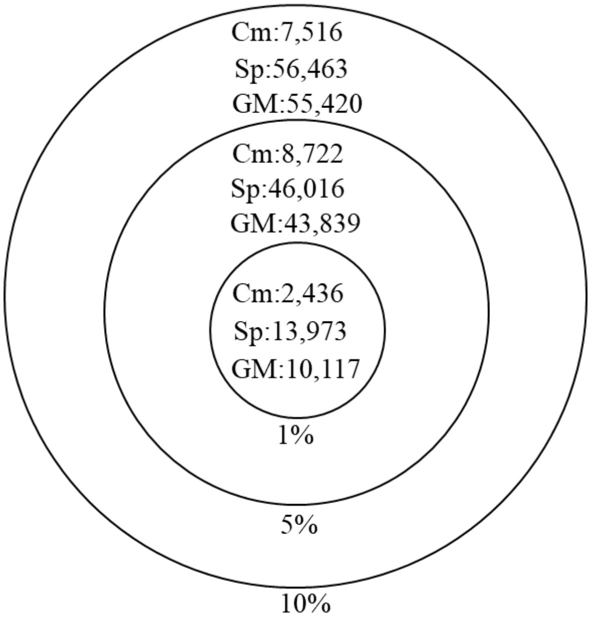


**Fig. S2**. The number of common and unique SNPs showing significant associations to Min.Tmp.Cld.M when and when not accounting for population structure. “Cm” depicts the number of SNPs identified by both Spearman (“Sp”) and GEMMA (“GM”) associations to climate. “Sp” depicts the number of significant associations identified by Spearman correlations only. “Gm” depicts the number of significant associations identified by GEMMA associations only. As thresholds for significance we used the $1^{st}$, $5^{th}$, and $10^{th}$ percentiles of the p-value distributions (Fig. S1). The numbers depicted at each threshold are mutually exclusive.

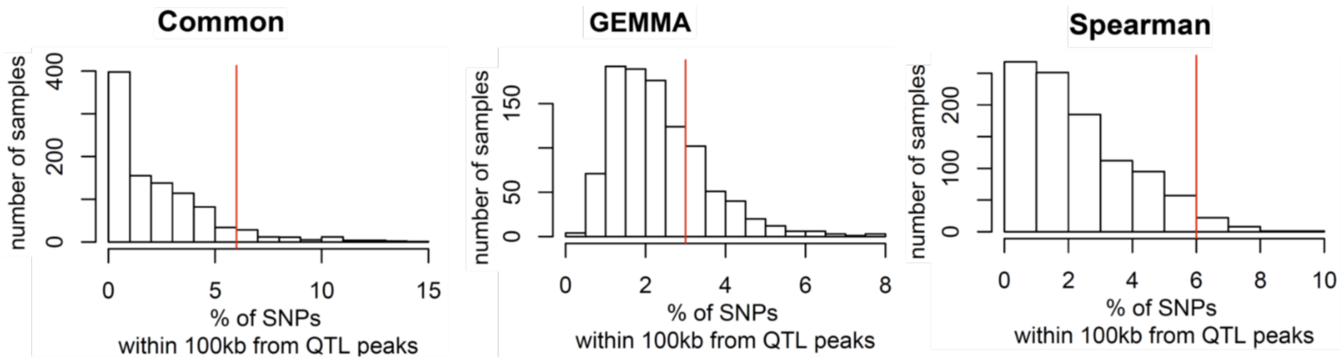


**Fig. S3**. The distribution of SNPs showing significant associations to Min.Tmp.Cld.M within 100kb from fitness QTL peaks. The observed percent (red line) of "Common" and "Spearman" SNPs was twice as high as the percent of "GEMMA" SNPs, and higher than the 95$^{th}$ percentiles of expected percentages derived using a circular permutation test.

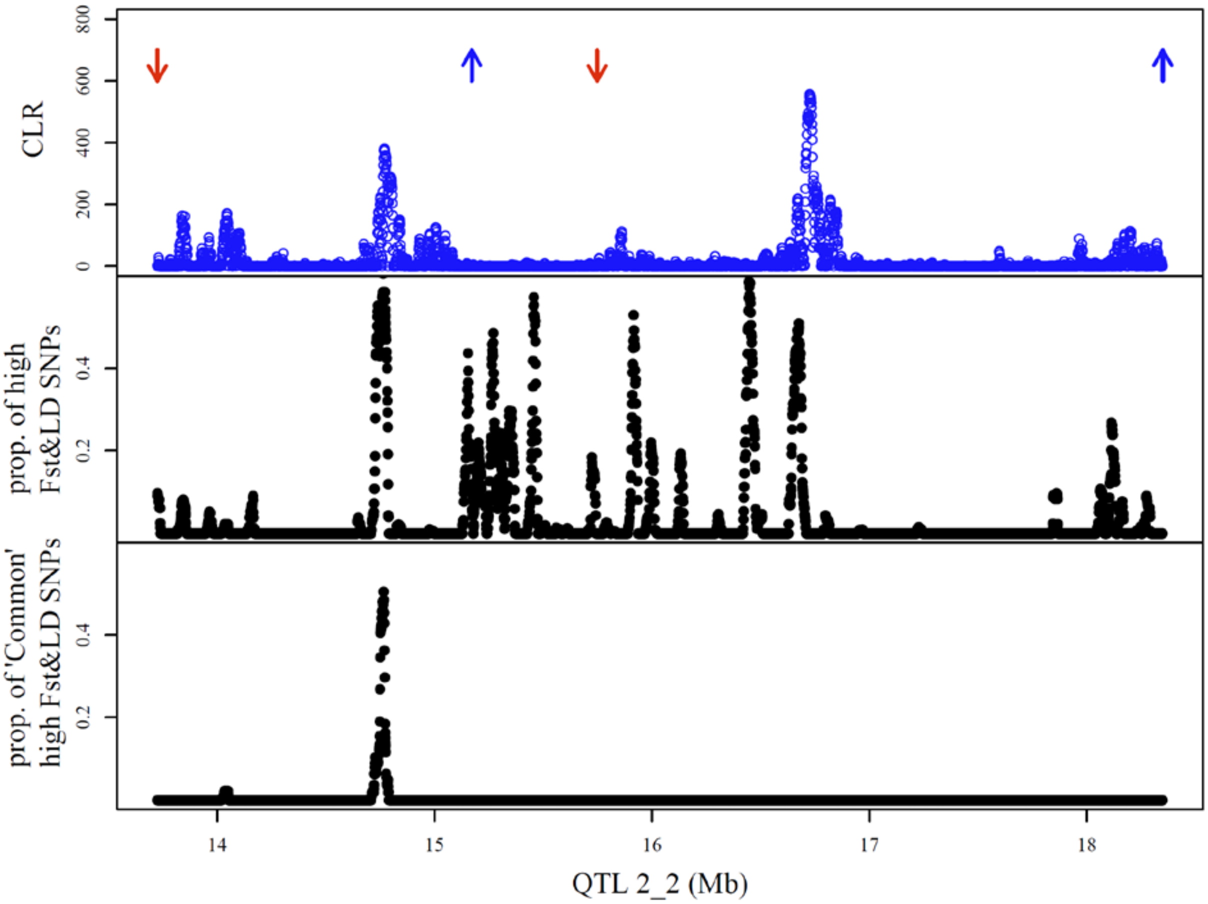


**Fig. S4.** Population genomic signatures of selection along genetic tradeoff QTL 2:2. The genetic tradeoff QTL was compiled using four fitness QTL (blue and red arrows-top panel). Blue arrows indicate locations where the Sweden genotype was associated with higher fitness than the Italy genotype in Sweden, and red arrows indicate locations where the Sweden genotype showed lower fitness than the Italy genotype in Italy. The top panel also depict Composite likelihood Ratios (CLR) for recent sweeps in North Sweden. Two regions showed CLRs >400, which are above the 99 percentilesof all CLR windows across the genome and above the neutral expectation (>85) previously identified (Price, et al. 2018) . The middle panel indicates the proportion of high $F_{ST}$ and LD SNPs within a 20kb window. The lower panel indicates the proportion of high $F_{ST}$ and LD SNPs that were also part of the set of SNPs that show significant associations to climate with and without accounting for population structure (‘Common’).

**Table S1.** Number of significant associations to four climate variables identified using GEMMA and using a FDR<0.1. False discovery rate was estimated using the ‘qvalue’ package implemented in R. The climate variables are Minimum Temperature of coldest Month (Min.Tmp.Cld.M). Photosynthetically active radiation during fall (PARFall) which is the time Italy and Sweden ecotypes germinate. Precipitation during warmest quarter of the year (Prec.Wrm.Q) and soil moisture (Soilm). Further details on how the climate variables were compiled are explained in (Lasky, et al. 2012)

| | FDR<0.1 |
|---|---:|
| Min.Tmp.Cld.M | 6,850 |
| PARFall | 146 |
| Prc.Wrm.Q. | 134 |
| Soilm | 519 |

**Table S2**. A list of 25 genes with cis-regulatory and/or nonsynonymous SNPs found in conserved regions and within 100 kb of fitness QTL peaks. The genes identified were close to fitness QTL that were part of three genetic tradeoff QTL (GT QTL), 2:2, 4:2, 5:5. The biological processes these genes were involved in were retrieved from the TAIR database.

| Gene | GT QTL | Biological process |
|---|---|---|
| AT2G36230 | 2:2 | histidine biosynthetic process, tryptophan biosynthetic process |
| AT2G44210 | 2:2 | Unknown |
| AT4G33140 | 4:2 | deoxyribonucleotide catabolic process |
| AT4G33150 | 4:2 | This is a splice variant of the LKR/SDH locus. It encodes a bifunctional polypeptide lysine-ketoglutarate reductase and saccharopine dehydrogenase involved in lysine degradation. There is another splice variant that encodes a mono saccharopine dehydrogenase protein. Gene expression is induced by abscisic acid, jasmonate, and under sucrose starvation. |
| AT4G33200 | 4:2 | actin filament organization, actin filament-based movement, nuclear migration,regulation of establishment or maintenance of cell polarity regulating cell shape |
| AT4G33240 | 4:2 | endomembrane system organization, phosphatidylinositol phosphorylation, pollen development, vacuole organization |
| AT4G33350 | 4:2 | protein folding, protein transport |
| AT4G33360 | 4:2 | farnesol metabolic process, negative regulation of abscisic acid-activated signaling pathway, terpenoid metabolic process |
| AT4G33380 | 4:2 | uknown |
| AT4G33410 | 4:2 | membrane protein proteolysis, signal peptide processing |
| AT4G33420 | 4:2 | hydrogen peroxide catabolic process, oxidation-reduction process, response to oxidative stress |
| AT4G33470 | 4:2 | histone deacetylation, tubulin deacetylation |
| AT5G64730 | 5:5 | unknown |
| AT5G64860 | 5:5 | unknown (DPE2 cold acclimation) |
| AT5G64930 | 5:5 | defense response, jasmonic acid and ethylene-dependent systemic resistance, jasmonic acid mediated signaling pathway, leaf senescence, photoperiodism, flowering, plant-type hypersensitive response, response to other organism, sugar mediated signaling pathway, systemic acquired resistance, trichome morphogenesis |
| AT5G65020 | 5:5 | phloem sucrose unloading, polysaccharide transport, primary root development,response to cold, response to heat, response to salt stress, response to water deprivation |
| AT5G65274 | 5:5 | uknown |
| AT5G65450 | 5:5 | protein deubiquitination, ubiquitin-dependent protein catabolic process |
| AT5G65460 | 5:5 | abscisic acid-activated signaling pathway, hyperosmotic salinity response,negative regulation of transcription, DNA-templated, response to chitin, response to cold |
| AT5G65530 | 5:5 | defense response to fungus, protein autophosphorylation, protein phosphorylation,trichome branching |
| AT5G65683 | 5:5 | gravitropism, root development |
| AT5G65690 | 5:5 | gluconeogenesis |
| AT5G66890 | 5:5 | defense response |
| AT5G66960 | 5:5 | proteolysis |
